# Dengue virus structural proteins are expressed on the surface of DENV-infected cells and are a target for antibody-dependent cellular phagocytosis

**DOI:** 10.1101/2024.07.21.604479

**Authors:** Mitchell J. Waldran, Elizabeth A. Kurtz, Chad J. Gebo, Timothy J. Rooney, Frank A. Middleton, Nathan H. Roy, Jeffrey R. Currier, Adam T. Waickman

## Abstract

Dengue virus (DENV) is an arboviral pathogen found in over 100 countries and a source of significant morbidity and mortality. While the mechanisms underpinning the pathophysiology of severe dengue are incompletely understood, it has been previously suggested that antibodies directed against the DENV envelope (E) protein can facilitate antibody dependent enhancement (ADE) of the infection, increasing the number of infected cells and the severity of infection in an exposed individual. Accordingly, there is interest in defining mechanisms for directly targeting DENV-infected cells for immunologic clearance, an approach that bypasses the risk of ADE. We have previously demonstrated that antibodies specific for DENV non-structural protein 1 (NS1) can opsonize and facilitate the phagocytic clearance of DENV infected cells. However, it is currently unclear if other DENV antigens are expressed on the surface of infected cells, and if these antigens can be targeted by antibody-dependent clearance mechanisms. In this study, we demonstrate that DENV structural proteins are expressed on the surface of DENV-infected cells, and that these antigens can be opsonized by both DENV-immune sera and monoclonal antibodies. In addition, DENV E-specific antibodies can facilitate phagocytic uptake of DENV-infected cells, resulting in the target-cell membrane localizing to endosomes of the engulfing phagocyte. Notably, no DENV genetic material was observed localizing to the engulfing phagocyte, suggesting that horizontal transfer of DENV from the infecting cell is an unlikely occurrence. In their totality, these data reinforce the concept that DENV E-reactive antibodies have a multifaceted role in DENV immunity and pathogenesis beyond neutralization and/or infection-enhancement.

## Introduction

Dengue virus (DENV) is a prevalent mosquito-borne pathogen that co-circulates in the tropics and sub-tropics as four antigenically and genetically distinct serotypes (DENV-1 to -4). Causing an estimated 100 million clinically apparent infections and 40,000 deaths every year, almost half of the world’s population is at risk of exposure to the virus due to the increasing range of its mosquito vectors *Aedes aegypti* and *Aedes albopictus* (1). DENV infections are typically characterized by fever, rashes, nausea, vomiting, and general aches and pains (1). However, a fraction of DENV-infected individuals progress to develop severe dengue, historically categorized as dengue hemorrhagic fever (DHF) or dengue shock syndrome (DSS). While the risk factors associated with developing severe disease are complex, the greatest single risk factor associated with progressing to severe dengue is a prior infection with a heterologous DENV serotype. The leading mechanistic explanation for this phenomenon is antibody dependent enhancement (ADE), wherein sub-neutralizing concentrations of DENV-specific IgG antibodies facilitate the uptake of DENV virions into FcγR expressing phagocytes during a secondary infection (2-6). This unique pathophysiological feature of dengue has complicated DENV countermeasure development for decades and has spurred interest in developing a more holistic understanding of the immunologic mechanisms associated with risk or protection from infection.

Much of the humoral immune response elicited by DENV infection is directed towards the structural components of the DENV virion; namely the envelope protein (E protein) and pre-membrane (prM) protein. Like all flaviviruses, the DENV E protein forms a homotypic dimer on the surface of a mature virion and is composed of three distinct domains: domain I, II, and III (7, 8). Domain I is primarily a structural center for the post fusion configuration of E, while domain II contains the fusion peptide that fuses to the viral envelope to the membrane of the endosome, thereby allowing for the release of the viral genome into the cytosol of the target cell (8-10). Domain III contains a putative receptor binding domain, facilitating interactions of the virion with DC-SIGN, mannose receptor, and other scavenger receptors used by DENV for cellular entry (7, 8).

After the resolution of an acute DENV infection, antibodies of multiple isotypes (IgG, IgM, and IgA) can be found in circulation that bind to prM, EDI, EDII, and EDIII, as well as non-structural protein 1 (NS1) (11, 12). Unlike the other non-structural DENV proteins, NS1 is expressed on the surface of – and secreted by – DENV infected cells, acting as a potent immunogen and convenient biomarker of acute DENV infection (12-14). Antibodies directed against the DENV envelope protein are responsible for facilitating virion neutralization, especially when directed against quaternary epitopes spanning multiple E dimers on the surface of the virion. However, IgG isotype antibodies targeting structural components of the DENV virion are also capable of facilitating ADE when present at sub-neutralizing concentrations (15, 16). Interestingly, IgA antibodies have not only been shown to not facilitate ADE, but they are capable of antagonizing IgG-mediated ADE (17, 18). This has inspired significant interest in understanding alternative antibody-dependent mechanisms for controlling DENV infection that do not require the direct interaction of antibodies with the DENV virion.

While virus-specific antibodies are most often defined by their neutralizing activity – meaning how well they can block virion/receptor interactions – many classes of antibodies also possess the ability to interact with Fc receptors expressed on leukocytes such as macrophages, monocytes, and NK cells. This antibody/FcR interaction can facilitate a broad range of Fc-dependent effector functions, including antibody-dependent cellular toxicity (ADCC) and antibody-dependent cellular phagocytosis (ADCP) (19). These effector mechanisms are thought to play a critical role in the clearance of virally infected cells, as viral antigens expressed on the surface of the infected cells allows for opsonization by virus-specific antibodies and subsequent phagocytic/lytic clearance by FcR expressing leukocytes. Our group has previously shown that DENV NS1 expressed on the surface of DENV-infected cells is readily opsonized by NS1-speific IgG and IgA isotype antibodies, thereby facilitating ADCP (20). However, NS1-specific antibodies are significantly less abundant in DENV-immune individuals than antibodies directed against DENV structural proteins, and prior evidence suggest that DENV structural proteins might also be expressed on the surface of infected cells (21). This leaves unaddressed the possibility that the abundant antibodies directed against directed against DENV structural proteins – such as prM and E – might also facilitate Fc-dependent effector mechanisms such as ACDP.

To fill this knowledge gap, we utilized an *in vitro* infection assay to quantify the expression of DENV proteins on the surface of DENV-infected cells. In addition to NS1 – which has previously been described to be expressed at high levels on the surface of infected cells - we observed surface expression of both prM and E, with EDIII-specific mAbs binding with particularly high efficiency. Antibodies directed against EDIII facilitated phagocytic uptake of cellular material from DENV-infected cells, resulting in this material localizing to lysosomes within the engulfing phagocyte. However, no transfer of DENV genomic material was detected during this process, suggesting that the risk of antibody-mediated horizontal transfer of infectious viruses is of low probability. Given the previously described association between EDIII-specific antibodies and protection from symptomatic DENV-infection, we posit that E-directed ADCP may be a mechanism for clearing virally-infected cells, paralleling processes previously described for NS1-specific antibodies.

## Results

### DENV-structural proteins are expressed on the surface of DENV infected cells

Our group – among others – has observed that DENV-immune sera can opsonize DENV infected cells, facilitating effector functions such as ADCP (20). The antigen that has been primarily cited to facilitate this interaction is NS1, which is known to be highly expressed on the surface of DENV-infected cells. To determine if DENV-immune serum contains opsonizing antibodies directed against DENV proteins other than NS1, we depleted NS1-specific antibodies from DENV immune serum. This was accomplished by repeated/sequential incubation of DENV-3 immune sera with a cell line engineered to express surface DENV NS1 (**Figure 1A**). The DENV-specific opsonizing activity of DENV naïve, DENV-immune serum (un-depleted), and anti-NS1-depleted serum was assessed using two cell lines: **1)** CEM.NK^R^ cells expressing DENV-3 NS1, **2)** DC-SIGN expressing CEM.NK^R^ cells infected with DENV-3. While the depletion of anti-NS1 antibodies completely abrogated opsonization of cells expressing only DENV NS1 (**Figure 1B, left**), the lack of NS1-specific antibodies only partially impacted the opsonization efficiency of DENV-infected cells by this same serum sample (**Figure 1B, right**). This result suggests that DENV proteins other than NS1 are found on the surface of infected cells and can serve as a target of antibody opsonization.

**Figure 1:**
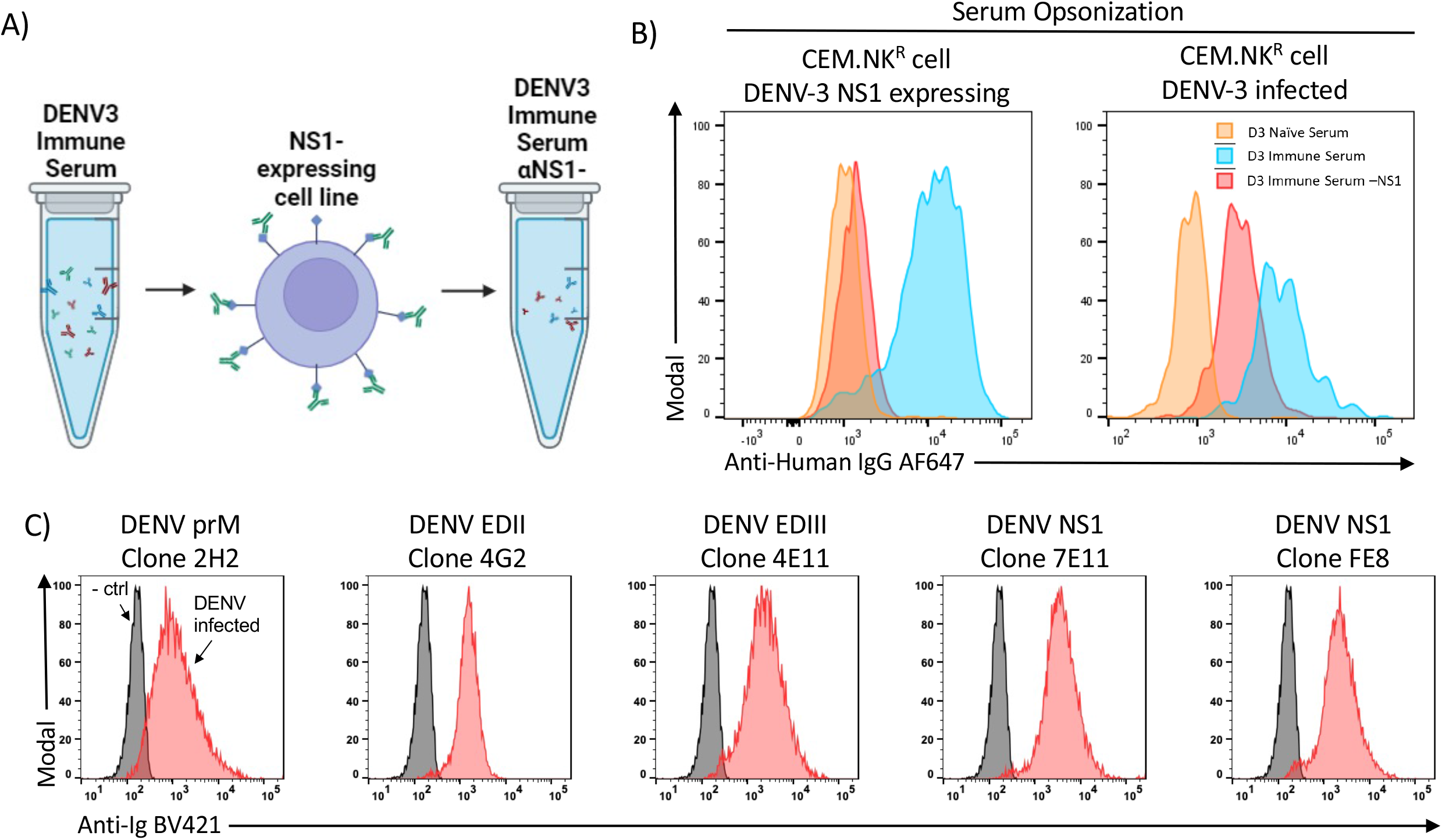
Identification of dengue virus proteins expressed on the surface of DENV-infected cells. **A)** Model for the depletion of NS1-reactive antibodies from whole DENV-immune serum as described in the methods **B)** Opsonization quantification on DENV3 NS1 expressing CEM.NK^R^ cells (left) and DENV3-infected CEM.NK^R^ cells (right) using either Naïve serum (orange), whole immune serum (blue) or whole serum that was depleted of NS1-reactive antibodies (red). Plots gated on cells positive for intracellular E antigen (4G2). **C)** Staining of control (grey) and DENV-3 infected (red) cells with the indicated monoclonal antibodies. DENV-3 infected samples gated on infected cells by intracellular anti-prM FITC staining.

To further define the composition of viral proteins on DENV-infected cells, we tested a well-characterized panel of mAbs specific for DENV viral proteins (**Figure 1C, Supplemental Figure 1**). As anticipated based on previously published work, NS1-specific antibodies readily bound to the surface of DENV-infected cells. However, in addition, we observed robust binding of prM-, EDII-, and EDIII specific mAbs to the surface of infected cells, with the EDIII-specific mAb 4E11 binding with notably high efficiency (**Figure 1C**), suggesting the antibodies directed against DENV structural proteins may have the potential Fc-dependent effector functions such as ADCP.

### DENV EDIII-specific antibodies can mediate monocytic phagocytosis of DENV3-infected cells

In light of the robust binding of EDIII specific antibodies described above, we wanted to investigate the ability of this class of antibody to facilitate Fc-dependent effector functions such as ADCP. To this end, we synthesized a pair of DENV EDIII-specific mAbs (clone 4E11) with either a human IgG1 or IgA1 Fc region. We confirmed that these isotype-switched antibodies retained their ability to bind DENV (**Figure 2A**) as well as opsonize DENV-infected cells (**Figure 2B**). To confirm the functional activity of these EDIII-specific mAbs, we first assessed the ability of these antibodies to mediate phagocytic-uptake of polystyrene beads coated with DENV E proteins. As predicted, both the IgG and IgA version of the EDIII-specific mAb facilitated the uptake of DENV-E conjugated polystyrene beads by primary human PBMC-derived monocytes **(Figure 2C**).

**Figure 2:**
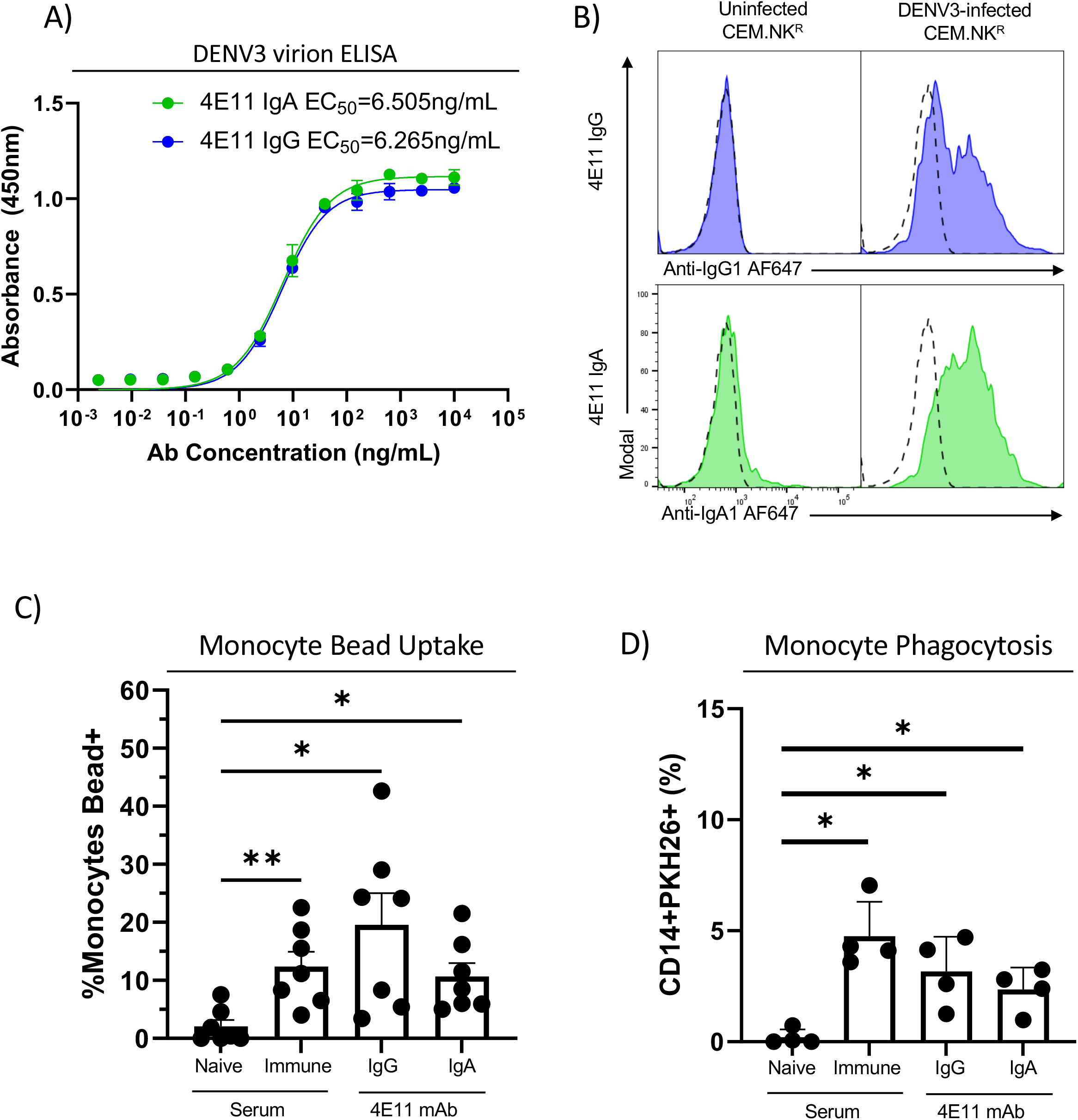
DENV envelope-specific IgG and IgA isotype monoclonal antibodies can mediate antibody-dependent cellular phagocytosis. **A)** Assessment of DENV E protein binding activity of 4E11 IgG and 4E11 IgA mAbs utilized in this study by ELISA, **B)** Opsonization of parental uninfected CEM.NK^R^ cells (left) and DENV3-infected CEM.NK^R^ cells (right), dashed lines indicating incubation without the primary 4E11 antibody, **C)** Percentage of monocytes that were positive for bead fluorescence, separated by antibody used to opsonize target. This data is expressed as background subtracted from a no serum condition, n=7 individual experiments. Error bars are mean ± SEM. * p < 0.05, paired one-way ANOVA, **D)** Percentage of monocytes that were positive for the target cell membrane dye, separated by antibody used to opsonize target. This data is expressed as background subtracted from a no serum condition, n=4 individual experiments. Error bars are mean ± SEM. * p < 0.05, paired one-way ANOVA

Having confirmed both the antigen-specificity and the functional activity of the 4E11 IgG and IgA mAbs, we next endeavored to determine the ability of these antibodies to mediate monocytic phagocytosis of DENV-infected cells. To this end, we utilized a previously described flow cytometry based ADCP assay (20). In brief, this assay begins by labeling the target cells, in this case the DENV3-infected CEM.NK^R^ cell line, with the lipophilic dye PKH26. These labeled target cells were then opsonized with DENV-naïve serum, DENV-immune serum, the aforementioned humanized 4E11 mAbs before being mixed with freshly isolated human PBMC. After an additional period of incubation, the heterogeneous sample is stained with anti-CD14 to allow for the identification of classical monocytes and the sample analyzed by flow cytometry. The percentage of CD14+ monocytes that are positive for the PKH26 dye is used as an indication of phagocytosis of the opsonized/PKH26-labeled infected cells. Using this assay, we observed that both DENV-immune serum and 4E11 IgG and IgA significantly increase the phagocytic ability of monocytes DENV3-infected cells **(Figure 2D)**. These results indicate that both IgG and IgA isotype antibodies directed against DIII of E protein can mediate phagocytic uptake of DENV-infected cells by primary human monocytes.

### Cellular material from opsonized DENV3-infected cells localizes to monocyte lysosomes

Having confirmed that EDIII-specific IgG and IgA mAbs can mediate phagocytic uptake of DENV-infected cells, we next wanted to assess the sub-cellular localization of engulphed material within the primary human monocytes utilized in this analysis. To investigate this, we followed the phagocytosis assay described above, and use flow cytometric sorting to isolate monocytes that phagocytized DENV-infected cell membrane. PBMC that were incubated in the absence of PKH26-labeled target cells were utilized as a negative control. These sorted cells were then labeled with Lysotracker Deep Red and imaged using epifluorescence microscopy.

Both control monocytes and monocytes positive for infected cellular membrane exhibited punctate lysosome staining (**Figure 3A**). In monocytes that contain material from the PKH26-labeled/DENV-infected target cells, the PKH26 signal also exhibited a punctate staining pattern that significantly co-localized with the lysosomal stain when quantified across multiple cells (**Figure 3A, Figure 3B**). These data suggest that material from DENV-infected cells localize to the endolysosomal pathway after engulfment by monocytes in an antibody-dependent fashion.

**Figure 3:**
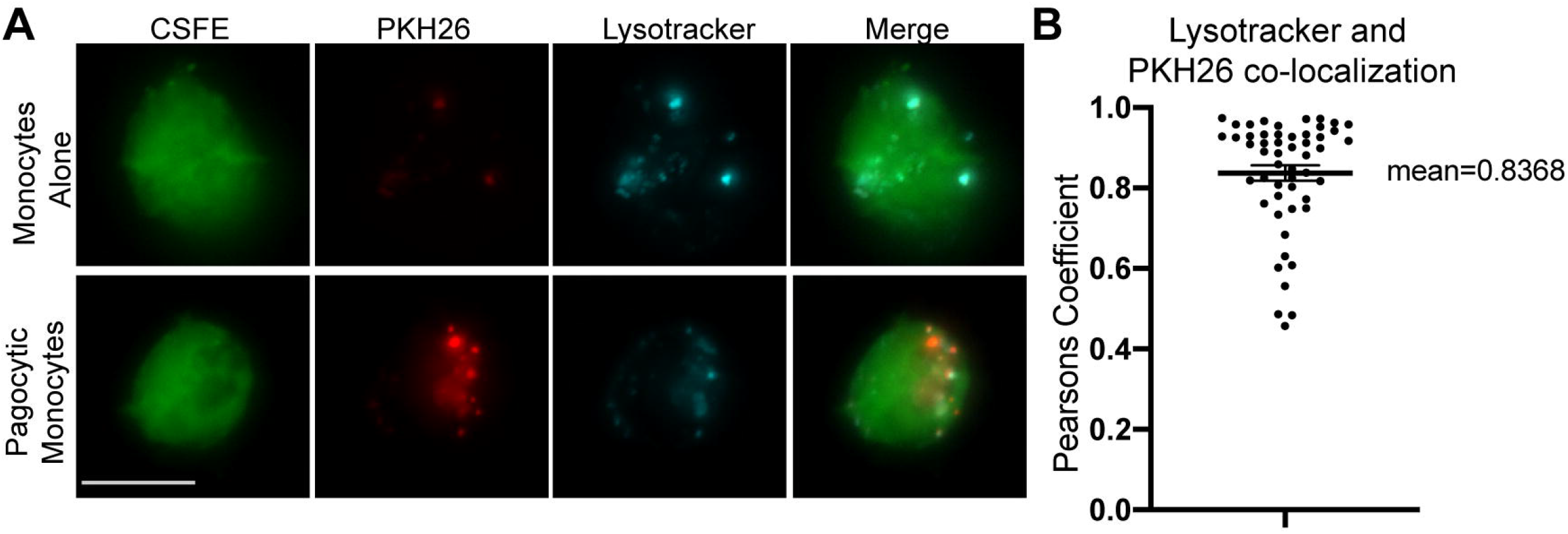
Intracellular trafficking of phagocytosed material from DENV-infected cells. **A)** Representative images of microscopy of monocytes that were incubated alone (top) or with DENV3 infected cells opsonized with human 4E11 IgG (bottom) at 63x magnification and each image was created using 3D projections with Z-stacks taken every 0.5um with nearest neighbors’ deconvolution. Monocyte cytosol was stained with CFSE (green), lysosomes were stained with Lysotracker Deep Red (cyan), and DENV3 infected cell membranes stained with PKH26 (red). Scale bar is 10 microns. Images were merged to show a visual representation of the colocalization, **B)** Quantification of colocalization between monocyte lysosomes and DENV3-infected cell membrane with each individual cell calculated separately, calculated using the Pearsons Coefficient from JACoP in FIJI, n=52

### Antibody-dependent phagocytosis does not lead to the horizontal transfer of DENV genomic material

A notable feature of the DENV life cycle is that the virus must localize to an acidified endosome to facilitate membrane fusion and escape into the cytoplasm. Considering the observation that EDIII-directed ADCP results in the localization of DENV-infected cellular material to a phagocytic lysosome, we wanted to determine if this process might facilitate the horizonal transfer of DENV genetic material from an infected cell to a susceptible phagocyte.

To investigate this potential route of DENV transfer, we performed the same ADCP assay described above and then utilized flow cytometry to isolate CD14+PKH26-monocytes (negative for DENV material), CD14+PKH26+ monocytes (positive for DENV material), and CD14-PKH26+ target cells (positive control for DENV genomic material) **(Figure 4A)**. Once the three populations were isolated, RNA was extracted and analyzed by high-sensitivity digital droplet PCR (ddPCR) for the presence of DENV-3 genomic RNA. The goal of this analysis was to compare abundance of DENV RNA in monocytes that were positive for material derived from DENV-infected cells and monocytes that were negative for material derived from DENV-infected cells, with the infected cells acting as the positive control for the presence of DENV RNA. While a robust DENV RNA signal was observed in the sorted target cell population, there was no difference in the abundance of DENV RNA detected in either sorted monocyte population. These results suggest that EDIII-mediated ADCP is not an efficient method for the horizontal transfer of DENV genomic material from infected cells to monocytes at the level of detection used in our ddPCR assay **(Figure 4B)**.

**Figure 4:**
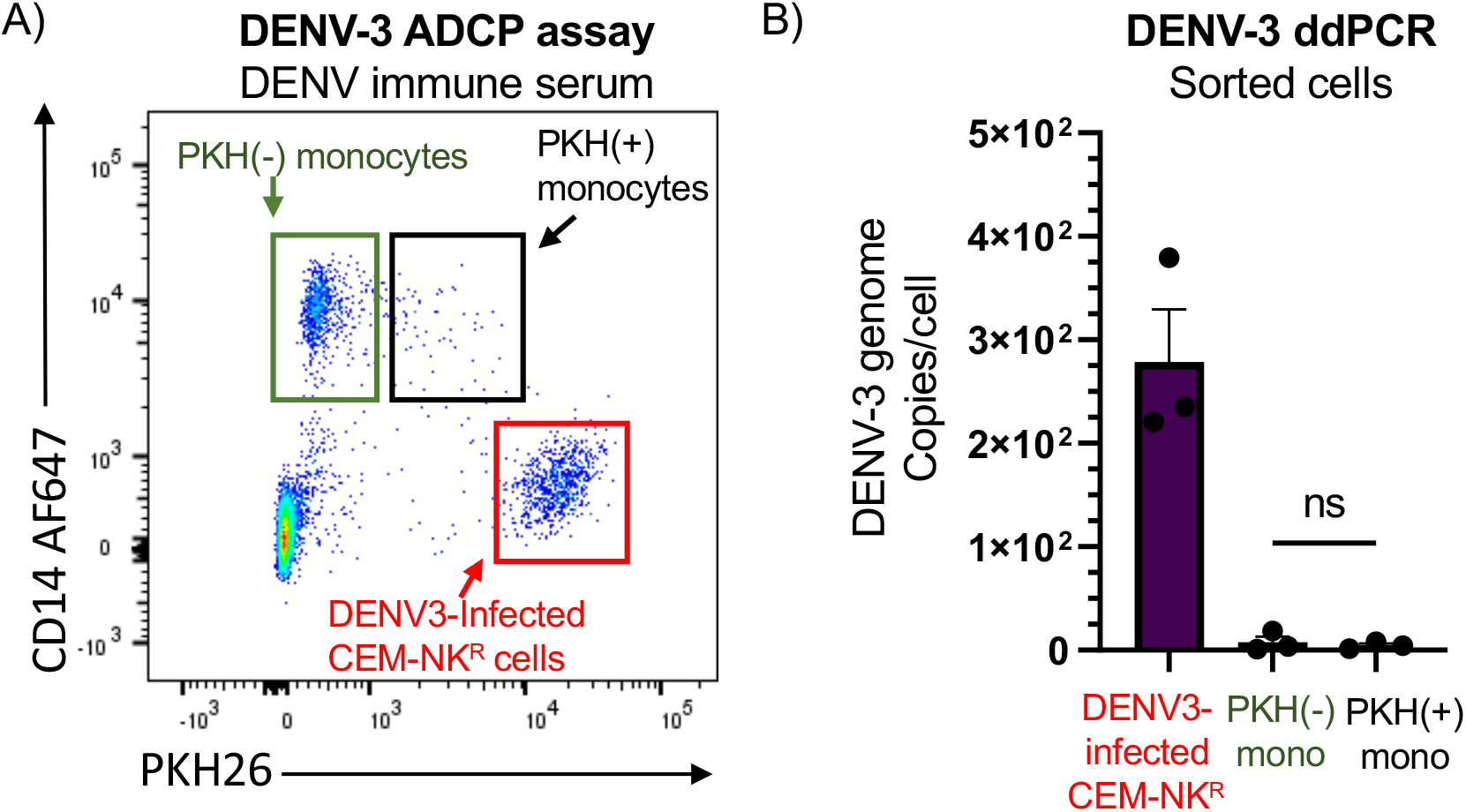
Quantification of DENV genomic transfer during antibody-mediated phagocytosis of DENV-infected cells. **A)** Example flow plot used to isolate out each cell type for use in the ddPCR. PKH-monocytes were taken from the single positive CD14+ cells, DENV3-infected target cells were taken from the single positive PKH26 membrane dye cells, and the PKH+ monocytes were taken from the double positive cells, **B)** Copies of DENV RNA per cell found using droplet digital PCR in DENV3-infected CEM.NK^R^ cells, PKH26(-) monocytes, and PKH26(+) monocytes that were incubated together and sorted separately using the Aria cell sorter. Error bars are mean ± SEM, paired one-way ANOVA, n=3

## Discussion

In this study, we describe a function for antibodies targeting DENV structural proteins in facilitating the phagocytosis of DENV-infected cellular material. This analysis demonstrated that DENV structural proteins are expressed on the surface of infected cells, and that both IgG and IgA antibodies recognizing DENV E protein can facilitate monocytic uptake of material from DENV-infected cells. In addition, we observed that the monocytic phagocytosis of DENV-infected cells leads to the internalization of infected cell material into lysosomes but did not result in the significant accumulation of DENV genomic material within the phagocyte.

While our study focuses on DENV, the generation of E-reactive antibodies after the resolution of infection is a common feature of mosquito-borne flavivirus infection in humans. This includes infections by Zika Virus, West Nile Virus, and Japanese Encephalitis Virus (22-24). Similar to DENV infection, antibodies targeting E protein also provide potent neutralizing functions to these pathogens (22-24). This study could herald a wider variety of therapeutics among DENV and similar viruses. For example, E-reactive antibodies, specifically IgA antibodies, could be used for more than just neutralizing infectious virions but also for the clearance of infected cells. The downside of using E-reactive antibodies is the risk of ADE (15-18). Our study shows that IgA and IgG E-reactive antibodies are equally productive in their ability to mediate phagocytosis. This is of note, as it has been shown that DENV-specific IgG is capable of mediating ADE, while IgA antibodies antagonize IgG-facilitated ADE (17, 18). This suggests the potential utility of an IgA-based therapeutic for DENV treatment with limited risk of disease enhancement (17, 18). In this light, the generation of E-reactive IgA antibodies may be a useful strategy to target not only virion neutralization, but also clearance of DENV-infected cells (17, 18).

There are a few limitations that should be considered while interpreting the results of this study. Most of the results in this study were generated using an immortalized cell line which has increased permissivity to DENV infection caused by the stable expression of DC-SIGN on the cell surface. The consequence of this is that the concentration of E protein in our assays might be significantly elevated compared to what is seen during natural infection, which may exaggerate the antibody-mediated effects seen for phagocytosis. Additional work is required to assess the *in vivo* contribution of E-directed ADCP in controlling DENV infection, and how this process may synergize with other antibody-dependent mechanisms of regulating DENV infection and pathogenesis. Regarding potential horizontal transfer of virus to effector cell, our results suggest that there is no significant difference in viral transfer between phagocytic and non-phagocytic monocytes. However, limitations of *in vitro* culture assays involving primary human monocytes make it difficult to conclusively demonstrate the lack of horizonal transfer of infectious virus via antibody-mediated phagocytosis. However, given the sensitive nature of the ddPCR assay utilized in this study we feel this is an unlikely scenario. Finally, we do not know the precise conformation and configuration of the E-protein on the surface of the infected cells. While it most likely not monomeric, the E-protein could exist as dimers such as on mature virions or perhaps even in the trimeric conformation that is present on immature virions (25) Hence, further work is required to fully elucidate the structure of E on the surface of DENV-infected cells.

In closing, we suggest that the results presented in this study show that DENV E-reactive antibodies can mediate more than just the neutralization of virions during a DENV-infection. Not only have we demonstrated a role for E-reactive antibodies outside of neutralization of virions, specifically IgA antibodies that do not cause ADE, but we also demonstrated that phagocytosis mediated by these antibodies did not increase horizontal transfer of virus from infected cell to effector cell. These findings provide valuable insight into the role of DENV E-reactive antibodies in DENV pathogenesis and clearance which may inform vaccine strategies and therapeutic development.

## Materials and Methods

### Cell lines and primary human PBMC isolation

A DENV-3 NS1 expressing cell line was generated by electroporating CEM.NK^R^ cells with a linearized plasmid containing the codon-optimized sequence were used for the serum depletion assays. NS1-expression was confirmed via flow cytometry. A DC-SIGN (CD209) expressing cell line was generated by transfecting CEM.NK^R^ cells with a linearized plasmid containing the codon-optimized sequence of DC-SIGN/CD209 (Genbank accession: NM_021155.4). Surface expression of DC-SIGN was confirmed by flow cytometry prior to each assay. PBMC were collected and isolated from normal healthy volunteers using BD Vacutainer CPT tubes containing sodium heparin. SUNY Upstate Medical University Institutional Review guidelines for the use of human subjects were followed for all experimental protocols in the study. All cell lines were cultured in R10 media (RPMI, 10% FBS, 1% Pen/Strep, and 1% L-glutamine).

### DENV-3 virion-capture ELISA

96 well NUNC MaxiSorb flat-bottom plates were coated with 2 μg/ml flavivirus group-reactive mouse monoclonal antibody 4G2 (Envigo Bioproducts, Inc.) diluted in borate saline buffer and incubated overnight at 4°C. The wells were washed 4 times with PBS-T and blocked with blocking buffer (0.25% BSA +1% Normal Goat Serum in PBS) for 30 minutes at RT. Blocking buffer was removed, and DENV-3 (strain CH53489) was captured for 2 hours then washed 4 times with PBS-T. 4E11 IgA and IgG monoclonal antibodies were prepared at 10ug/mL top concentration followed by 4-fold serial dilutions, plated, and incubated for 2 hours at RT. The plates were washed 4 times with PBS-T. Peroxidase-labeled anti-human IgG (Southern Biotech 2044-05) or IgA (Biolegend 411002) were added to the wells and incubated for 1 hour at RT. The plates were washed 4 times with PBS-T, and TMB substrate was added to the wells for 10 minutes and read on a Biotek μQuant plate reader at 450nm.

### NS1-reactive Antibody Depletion

10^7^ DENV3-NS1-expressing CEM.NK^R^ cells were incubated with serum diluted 1:50 in FACs buffer from patients experimentally infected with DENV3 for 1 hour at 4 □C on a gentle rocker in a 15mL conical tube. The cells were then centrifuged for 5 minutes at 500g, and the serum was collected and used to resuspend another tube full of 10^7^ DENV3-NS1-expressing CEM.NK^R^ cells, with the first tube being thrown away. This process was repeated, with regular verification of the depletion step by staining the incubated cells with an anti-human IgG fluorescent antibody and running these cells on the LSRII flow cytometer (BD Biosciences) **(Figure 1A)**. Once the verification showed no opsonization of NS1-expressing cells by the depleted serum, the depleted serum was stored at 4 □C.

### Antibody-Dependent Phagocytosis Bead Assay

Recombinant DENV3 E protein (abcam, AB256438-1001) was biotinylated using the EZ-LINK Sulfo-NHS-LC biotinylation kit (Life Technologies, 21435). The biotinylated E-protein was then incubated with 1um fluospheres bound with NetrAvidin (Invitrogen F8776) overnight at 4 □C. The next day, the beads were loaded into the appropriate wells of a polypropylene round-bottom 96 well plate at 10^6 beads/well and washed twice with PBS + 2% FBS. They were then incubated with sera (1:500 dilution) or anti-E monoclonal antibodies (mAbs) (1μg/mL) for 30 minutes at room temperature. Purified PBMC (2×10^5^ cells per well) was directly added to the antibody and bead mixture and incubated for 30 minutes at 37°C. The wells were then washed twice with PBS + 2% FBS and then stained with anti-human CD14 (Biolegend 301818 clone M5E2) for 20 minutes at room temperature, washed again, and analyzed on the LSRII flow cytometer (BD Biosciences).

### Antibody-Dependent Phagocytosis Assay

CEM.NK^R^ and DENV3 infected DC-SIGN-expressing CEM.NK^R^ cells were stained with PKH26 at a final concentration of 3.0×10^−6^M resuspended in Diluent C for 5 minutes at room temperature. Cells were then washed twice with PBS + 2% FBS prior to opsonization at 4°C with sera (1:500 dilution) or anti-E mAbs (1μg/mL) for 30 minutes. After opsonization, 2×10^4^ of the labeled CEM.NK^R^ cells were added to a well of a polypropylene round-bottom 96 well plate and mixed with 2×10^5^ of purified PBMC at 37°C. The cells were incubated together for 3 hours at 37 □C. The cells were then washed twice, stained with anti-human CD14 (Biolegend 301818 clone M5E2) for 20 minutes at room temperature, washed again, and analyzed on the LSRII flow cytometer (BD Biosciences).

### Microscopy

The above ADCP assay was performed; however, the monocytes were stained with an anti-CD14 BV510 (Biolegend, 367123) antibody then sorted out using the Aria Cell Sorter. Monocytes were then stained with CFSE (BD Horizon, 565082) and Lysotracker Deep Red (Invitrogen, L12492) before being transferred to an 8-well microscope plate (ibidi 80806). Cells were allowed to settle for 15 minutes before being imaged on a Marianas system (3i) enclosed in an environmental chamber (Okolab) consisting of a Ziess Axio Observer 7 equipped with a X-Cite mini+ light source (Excelitas) and a Prime BSI Express CMOS camera (Photometrics). Images were taken at 63x magnification with each image created using 3D projections with Z-stacks taken every 0.5um with nearest neighbors’ deconvolution using SlideBook 6 (3i) done before exporting to Fiji for analysis (26).

### Colocalization Quantification

The images from the above microscopy were scaled normalizing each channel (target cell membrane stain, monocyte marking stain, and lysosomal stain). Scaled channels were then merged and cropped for analysis of individual cells. Cells were then analyzed using the Fiji plugin, JACoP, and quantified for overlap between the target cell membrane stain and the lysosomal dye using Pearsons’s coefficient. The results are reported as the Pearson’s coefficient for the overlap of each individual cell.

### DENV-3 Digital ddPCR

The ADCP assay described above was followed using the DENV 3 infected cells and 4E11 IgG mAb condition. Instead of being analyzed using a flow cytometer, the cells were isolated using the Aria Cell Sorter. The DENV3 infected CEM.NK^R^ cells, target cell membrane negative monocytes, and target cell membrane positive cells were each isolated from the ADCP mixture.

DENV3 infected cell samples underwent RNA purification by first passing them through QIAshredder (Qiagen Catalog #79656) columns to rupture cell membranes and disrupt other cellular structures. This procedure was conducted by following the manufacturer’s instructions. DENV3 RNA was then purified from the shredded samples with RNeasy Mini Kits (Qiagen Catalog #74104) by adhering to the manufacturer’s instructions. The eluted RNA samples were stored in a -80 **°**C freezer for no longer than two months before being used in ddPCR experiments.

Primers and probe targeting the gene of DENV3 capsid protein were used with the One-Step-ddPCR Advanced Kit for Probes (Bio-Rad Catalog #1864022). The oligonucleotides were manufactured by Integrated DNA Technologies (the oligonucleotide sequences are available upon request). For one reaction, 6.25 uL of Supermix, 2.5 uL of reverse transcriptase, 1.25 uL of 300 mM Dithiothreitol, 1.2 uL of primer/probe mix, 5 uL of RNA, and 8.8 uL of water was combined to make a 25 uL solution. The primer/probe mix had forward and reverse primers at a concentration of 12.5 uM and probe at 5 uM. This gives a final concentration in the PCR solution of 0.625 uM for the primers and 0.25 uM for the probe. 20 uL of the reaction mixture was used for droplet generation done by the QX200 Droplet Generator (Bio-Rad Catalog # 1864003). The plate was sealed by a PX1 PCR Plate Sealer (Bio-Rad Catalog #1814000). Reverse transcription PCR was then performed using a C1000 Touch Thermal Cycler (Bio Rad Catalog #1851196) under the following conditions: 50 °C for 60 mins, 95 °C for 10 mins, followed by 44 cycles of 95 °C for 30 seconds (DNA melting) and 59.8 °C for 1 min with a 2 °C/s ramp rate (annealing), 98 °C for 10 mins, and finally 4 °C for 30 mins. Droplet fluorescence in the FAM channel was measured by a QX200 Droplet Reader (Bio-Rad Catalog #1864003). The threshold was determined automatically by the QuantaSoft software (Version 1.7.4.0917), and the plots were visually examined to verify the threshold setting and the quality of the data. The software provides the DNA concentration of the reaction volume. To determine the concentration of the input sample, the measured concentration was multiplied by the ratio of the reaction volume (25 uL) to the RNA sample volume.

Sensitivity and specificity testing were done to validate the assay. Purified DENV3 RNA (strain CH53489, GenBank ID: DQ863638.1) was serially diluted to determine the linear range of the ddPCR assay and to measure the concentration of the purified RNA starting material. The Limit of Quantification (LoQ) was assessed by diluting the purified RNA 1000, 4000, and 6000-fold and then measuring the target DNA concentration of replicate samples (n=5) by RT-ddPCR. An LoQ with a CV of 25% or lower was considered acceptable. The 6000-fold diluted replicates yielded an average of 14.75 copies/uL with a CV of 20.16%. Specificity testing involved determining whether the DENV3 assay would detect different serotypes of dengue virus (DENV1, 2, and 4). Cell-culture supernatant samples of these three other serotypes of dengue were tested in triplicate using the DENV3 assay by RT-ddPCR and there was no positive droplets were seen in any of the nine reactions, indicating complete target specificity through 44 cycles of PCR. DENV3 cell-culture supernatant samples consistently were amplified and detected by RT-ddPCR.

### Opsonization assays

DENV3 infected CEM.NK^R^ cells were incubated with serum or monoclonal antibody, depending on the assay, for 30 minutes at room temperature in PBS + 2% FBS. The cells were then washed twice, and then incubated with fluorescently tagged anti-human IgG (Southern Biotech, 2040-31) or IgA (Southern Biotech, 2050-31) antibodies for 30 minutes at room temperature. The cells were then washed two more times before being analyzed on the LSRII flow cytometer.

### Statistical analysis

All statistical analyses were performed using GraphPad Prism Software (GraphPad Software, La Jolla, CA). A P-value <0.05 was considered significant. ANOVA was used for the comparison of multiple conditions.

## ACKNOWLEDGEMENTS

We gratefully acknowledge the excellent technical assistance provided by Lisa Phelps of the SUNY Upstate Medical University Flow Cytometry Core. We also acknowledge Céline S.C. Hardy of the Waickman lab for her contributions to editing and reviewing the manuscript.

## Author contributions

**Conceptualization:** M.J.W., J.R.C., A.T.W., **Formal analysis:** M.J.W., A.T.W. **Funding acquisition:** A.T.W. **Investigation:** M.J.W., E.A.K., C.J.G., T.J.R. **Writing – original draft:** M.J.W., A.T.W. **Writing – review and editing:** all authors

## Disclaimer

Material has been reviewed by the Walter Reed Army Institute of Research. There is no objection to its presentation and/or publication. The opinions or assertions contained herein are the private views of the authors, and are not to be construed as official, or as reflecting true views of the National Institutes of Health, the Department of the Army or the Department of Defense. The investigators have adhered to the policies for protection of human subjects as prescribed in AR 70–25.

## References

1. Bhatt, S., P. W. Gething, O. J. Brady, J. P. Messina, A. W. Farlow, C. L. Moyes, J. M. Drake, J. S. Brownstein, A. G. Hoen, O. Sankoh, M. F. Myers, D. B. George, T. Jaenisch, G. R. Wint, C. P. Simmons, T. W. Scott, J. J. Farrar, and S. I. Hay. 2013. The global distribution and burden of dengue. Nature 496: 504–507.

2. Halstead, S. B. 1979. In vivo enhancement of dengue virus infection in rhesus monkeys by passively transferred antibody. J Infect Dis 140: 527–533.

3. Halstead, S. B. 2014. Dengue Antibody-Dependent Enhancement: Knowns and Unknowns. Microbiol Spectr 2.

4. Halstead, S. B., and E. J. O’Rourke. 1977. Dengue viruses and mononuclear phagocytes. I. Infection enhancement by non-neutralizing antibody. J Exp Med 146: 201–217.

5. Halstead, S. B., H. Shotwell, and J. Casals. 1973. Studies on the pathogenesis of dengue infection in monkeys. I. Clinical laboratory responses to primary infection. J Infect Dis 128: 7–14.

6. Halstead, S. B., H. Shotwell, and J. Casals. 1973. Studies on the pathogenesis of dengue infection in monkeys. II. Clinical laboratory responses to heterologous infection. J Infect Dis 128: 15–22.

7. Costin, J. M., E. Jenwitheesuk, S. M. Lok, E. Hunsperger, K. A. Conrads, K. A. Fontaine, C. R. Rees, M. G. Rossmann, S. Isern, R. Samudrala, and S. F. Michael. 2010. Structural optimization and de novo design of dengue virus entry inhibitory peptides. PLoS Negl Trop Dis 4: e721.

8. Modis, Y., S. Ogata, D. Clements, and S. C. Harrison. 2004. Structure of the dengue virus envelope protein after membrane fusion. Nature 427: 313–319.

9. Chambers, T. J., C. S. Hahn, R. Galler, and C. M. Rice. 1990. Flavivirus genome organization, expression, and replication. Annu Rev Microbiol 44: 649–688.

10. Uno, N., and T. M. Ross. 2018. Dengue virus and the host innate immune response. Emerg Microbes Infect 7: 167.

11. Rothman, A. L. 2004. Dengue: defining protective versus pathologic immunity. J Clin Invest 113: 946–951.

12. Waickman, A. T., J. Q. Lu, H. Fang, M. J. Waldran, C. Gebo, J. R. Currier, L. Ware, L. Van Wesenbeeck, N. Verpoorten, O. Lenz, L. Tambuyzer, G. Herrera-Taracena, M. Van Loock, T. P. Endy, and S. J. Thomas. 2022. Evolution of inflammation and immunity in a dengue virus 1 human infection model. Sci Transl Med 14: eabo5019.

13. Lyke, K. E., J. V. Chua, M. Koren, H. Friberg, G. D. Gromowski, R. R. Rapaka, A. T. Waickman, S. Joshi, K. Strauss, M. K. McCracken, H. Gutierrez-Barbosa, B. Shrestha, C. Culbertson, P. Bernal, R. A. De La Barrera, J. R. Currier, R. G. Jarman, and R. Edelman. 2024. Efficacy and immunogenicity following dengue virus-1 human challenge after a tetravalent prime-boost dengue vaccine regimen: an open-label, phase 1 trial. Lancet Infect Dis.

14. Waickman, A. T., K. Newell, J. Q. Lu, H. Fang, M. Waldran, C. Gebo, J. R. Currier, H. Friberg, R. G. Jarman, M. D. Klick, L. A. Ware, T. P. Endy, and S. J. Thomas. 2024. Low-dose dengue virus 3 human challenge model: a phase 1 open-label study. Nat Microbiol 9: 1356–1367.

15. de Alwis, R., K. L. Williams, M. A. Schmid, C. Y. Lai, B. Patel, S. A. Smith, J. E. Crowe, W. K. Wang, E. Harris, and A. M. de Silva. 2014. Dengue viruses are enhanced by distinct populations of serotype cross-reactive antibodies in human immune sera. PLoS Pathog 10: e1004386.

16. Williams, K. L., S. Sukupolvi-Petty, M. Beltramello, S. Johnson, F. Sallusto, A. Lanzavecchia, M. S. Diamond, and E. Harris. 2013. Therapeutic efficacy of antibodies lacking Fcgamma receptor binding against lethal dengue virus infection is due to neutralizing potency and blocking of enhancing antibodies [corrected]. PLoS Pathog 9: e1003157.

17. Wegman, A. D., H. Fang, A. L. Rothman, S. J. Thomas, T. P. Endy, M. K. McCracken, J. R. Currier, H. Friberg, G. D. Gromowski, and A. T. Waickman. 2021. Monomeric IgA Antagonizes IgG-Mediated Enhancement of DENV Infection. Front Immunol 12: 777672.

18. Wegman, A. D., M. J. Waldran, L. E. Bahr, J. Q. Lu, K. E. Baxter, S. J. Thomas, and A. T. Waickman. 2023. DENV-specific IgA contributes protective and non-pathologic function during antibody-dependent enhancement of DENV infection. PLoS Pathog 19: e1011616.

19. Forthal, D. N. 2014. Functions of Antibodies. Microbiol Spectr 2: 1–17.

20. Waldran, M. J., A. D. Wegman, L. E. Bahr, N. H. Roy, J. R. Currier, and A. T. Waickman. 2023. Soluble NS1 Antagonizes IgG- and IgA-Mediated Monocytic Phagocytosis of DENV Infected Cells. J Infect Dis 228: 70–79.

21. Aisenberg, L. K., K. E. Rousseau, K. Cascino, G. Massaccesi, W. H. Aisenberg, W. Luo, K. Muthumani, D. B. Weiner, S. S. Whitehead, M. A. Chattergoon, A. P. Durbin, and A. L. Cox. 2022. Cross-reactive antibodies facilitate innate sensing of dengue and Zika viruses. JCI Insight 7.

22. Zaidi, M. B., L. Cedillo-Barron, Y. A. M. E. Gonzalez, J. Garcia-Cordero, F. D. Campos, K. Namorado-Tonix, and F. Perez. 2020. Serological tests reveal significant cross-reactive human antibody responses to Zika and Dengue viruses in the Mexican population. Acta Trop 201: 105201.

23. Goo, L., K. Debbink, N. Kose, G. Sapparapu, M. P. Doyle, A. W. Wessel, J. M. Richner, K. E. Burgomaster, B. C. Larman, K. A. Dowd, M. S. Diamond, J. E. Crowe, Jr., and T. C. Pierson. 2019. A protective human monoclonal antibody targeting the West Nile virus E protein preferentially recognizes mature virions. Nat Microbiol 4: 71–77.

24. Goncalvez, A. P., C. H. Chien, K. Tubthong, I. Gorshkova, C. Roll, O. Donau, P. Schuck, S. Yoksan, S. D. Wang, R. H. Purcell, and C. J. Lai. 2008. Humanized monoclonal antibodies derived from chimpanzee Fabs protect against Japanese encephalitis virus in vitro and in vivo. J Virol 82: 7009–7021.

25. Kostyuchenko, V. A., Q. Zhang, J. L. Tan, T. S. Ng, and S. M. Lok. 2013. Immature and mature dengue serotype 1 virus structures provide insight into the maturation process. J Virol 87: 7700–7707.

26. Schindelin, J., I. Arganda-Carreras, E. Frise, V. Kaynig, M. Longair, T. Pietzsch, S. Preibisch, C. Rueden, S. Saalfeld, B. Schmid, J. Y. Tinevez, D. J. White, V. Hartenstein, K. Eliceiri, P. Tomancak, and A. Cardona. 2012. Fiji: an open-source platform for biological-image analysis. Nat Methods 9: 676–682.

