## Supplemental Figures for "Dengue virus structural proteins are expressed on the surface of DENV-infected cells and are a target for antibody-dependent cellular phagocytosis"

Supplemental 1

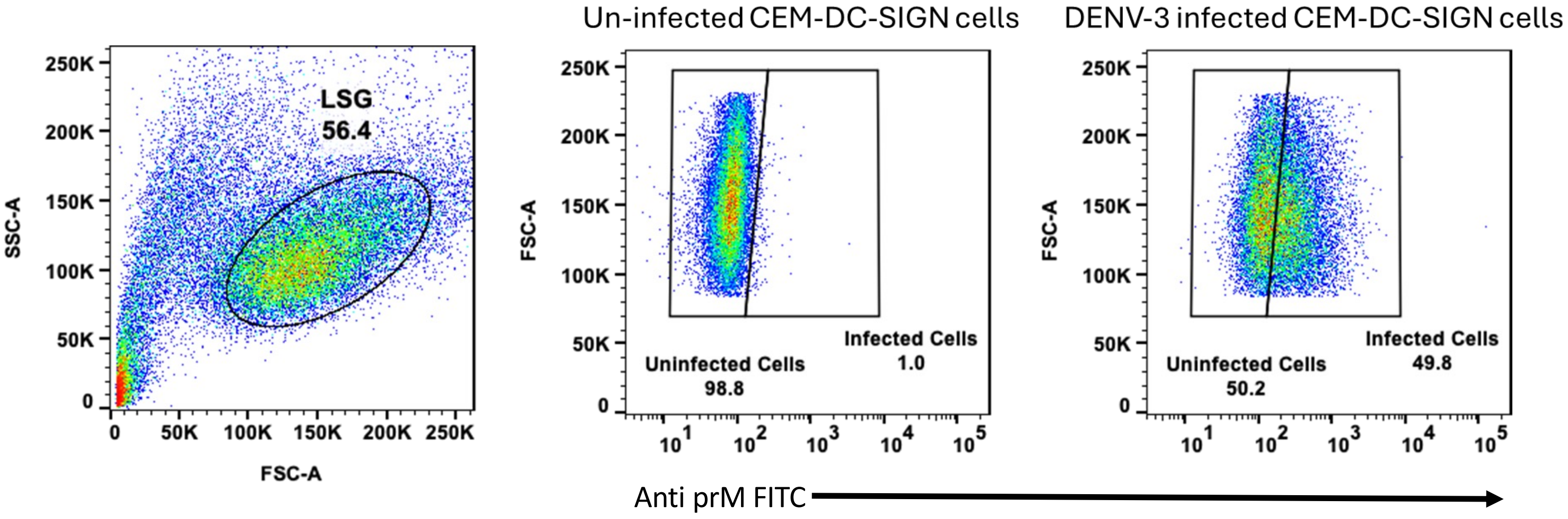


**Supplemental Figure 1:** Gating strategy used in the staining of DENV3-infected cells with 2H2, 4G2, 4E11, 7E11, and FE8. Shown are the gates before the reported graphs in Figure 1C.

Supplemental 2

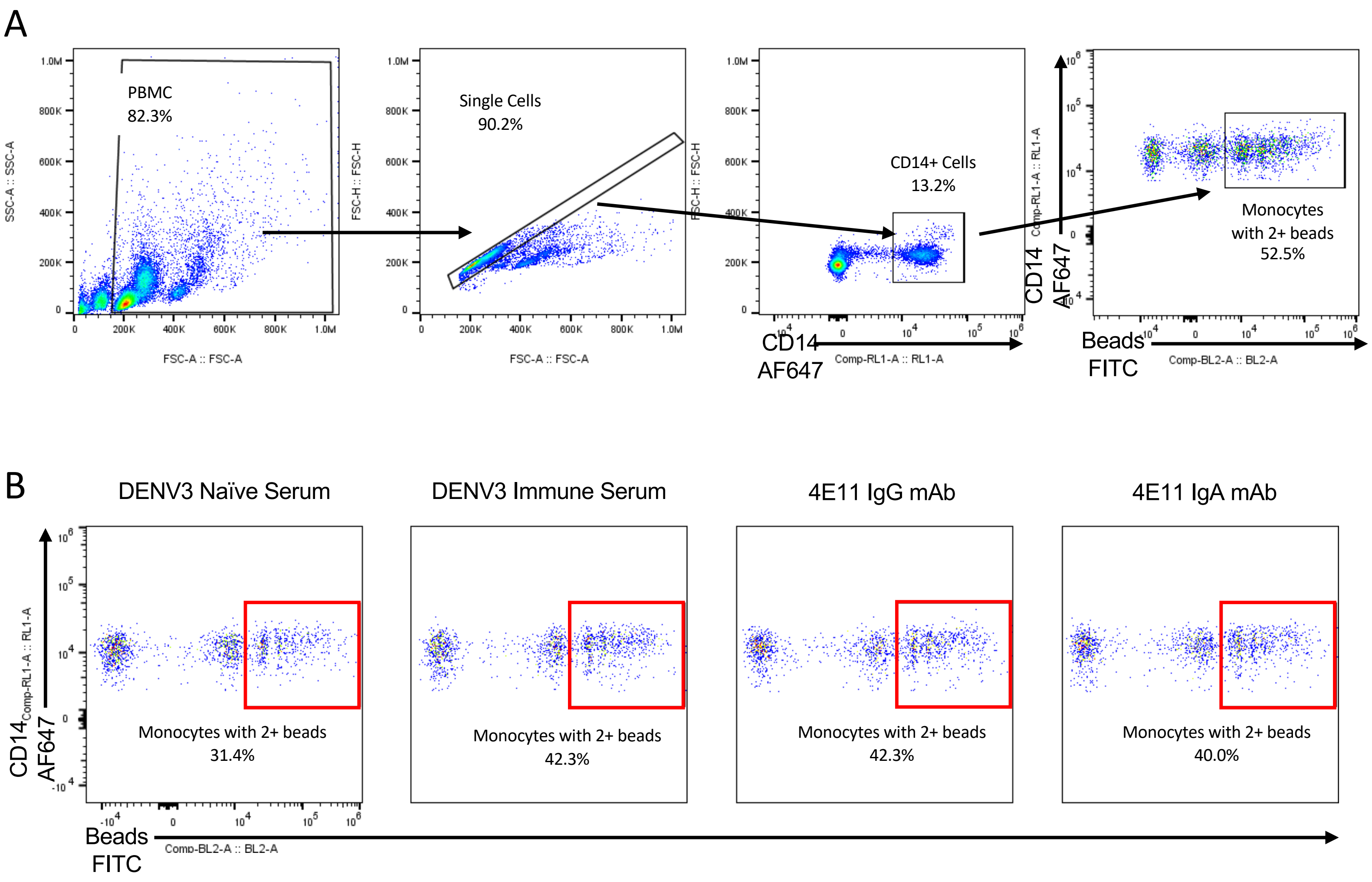


**Supplemental Figure 2: Gating strategy and example flow plots for Bead phagocytosis:** A) Gating strategy for bead phagocytosis assay, B) Representative flow plots from the bead phagocytosis assay.

Supplemental 3

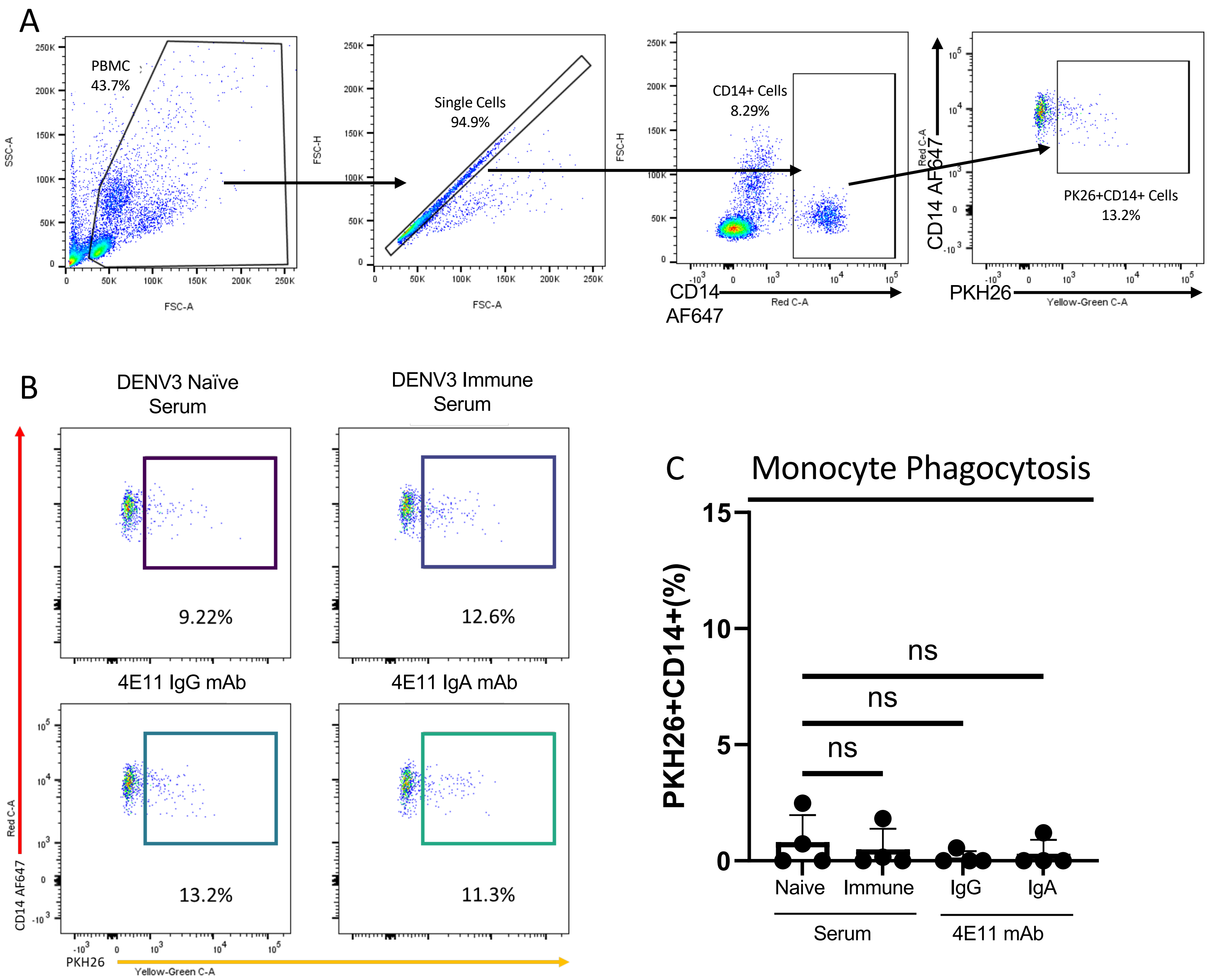


**Supplemental Figure 3: Gating strategy and example flow plots for DENV3-infected cell phagocytosis:** A) Gating strategy for DENV3-infected cell phagocytosis assay, B) Representative flow plots from the DENV3-infected cell phagocytosis assay, C) Monocyte phagocytosis of uninfected parental cells with the percentage of monocytes that were positive for the target cell membrane dye, separated by antibody used to opsonize target. This data is expressed as background subtracted from a no serum condition, n=4 individual experiments. Error bars are mean ± SEM. * $p < 0.05$, paired one-way ANOVA
